# A robust voice-selective response in the human brain as revealed by electrophysiological recordings and fast periodic auditory stimulation

**DOI:** 10.1101/2021.03.13.435142

**Authors:** Francesca M. Barbero, Roberta P. Calce, Siddharth Talwar, Bruno Rossion, Olivier Collignon

## Abstract

Voices are arguably among the most relevant sounds in humans’ everyday life, and several studies have suggested the existence of voice-selective regions in the human brain. Despite two decades of research, defining the human brain regions supporting voice recognition remains challenging. Moreover, whether neural selectivity to voices is merely driven by acoustic properties specific to human voices (e.g. spectrogram, harmonicity), or whether it also reflects a higher-level categorization response is still under debate. Here, we objectively measured rapid automatic categorization responses to human voices with Fast Periodic Auditory Stimulation (FPAS) combined with electroencephalography (EEG). Participants were tested with stimulation sequences containing heterogeneous non-vocal sounds from different categories presented at 4 Hz (i.e., 4 stimuli/second), with vocal sounds appearing every 3 stimuli (1.333 Hz). A few minutes of stimulation are sufficient to elicit robust 1.333 Hz voice-selective focal brain responses over superior temporal regions of individual participants. This response is virtually absent for sequences using frequency-scrambled sounds, but is clearly observed when voices are presented among sounds from musical instruments matched for pitch and harmonicity-to-noise ratio. Overall, our FPAS paradigm demonstrates that the human brain seamlessly categorizes human voices when compared to other sounds including matched musical instruments and that voice-selective responses are at least partially independent from low-level acoustic features, making it a powerful and versatile tool to understand human auditory categorization in general.

**Significance statement:** Voices are arguably among the most relevant sounds we hear in our everyday life, and several studies have corroborated the existence of regions in the human brain that respond preferentially to voices. However, whether this preference is driven by specific acoustic properties of voices or if it rather reflects a higher-level categorization response to voices is still under debate. We propose a new approach to objectively identify rapid automatic voice-selective responses with frequency tagging and electroencephalographic recordings. In four minutes of recording only, we recorded robust voice-selective responses independent from low-level acoustic cues, making this approach highly promising for studying auditory perception in children and clinical populations.

## 1. Introduction

Voices of conspecifics are arguably among the most relevant sounds we hear in our everyday life: they do not only carry speech, but also convey a wealth of information about the speakers such as their sex, age, emotional status, identity, trustworthiness, etc. (Belin et al., 2004). Regions in the human brain located along the bilateral superior temporal sulcus (STS) respond more to human voices than to other sounds (“Temporal Voice Areas”, TVAs), playing a key role in voice recognition (Belin et al., 2002, 2000). However, whether this selectivity is fully accounted for by specific acoustic properties of voices (Moerel et al., 2012; Ogg et al., 2019; Staeren et al., 2009), or if it also reflects a higher-level categorization response beyond these low-level auditory properties is still under debate. This question has been previously addressed through careful choice and design of acoustic stimuli (Agus et al., 2017; Belin et al., 2002; Levy et al., 2003) and sophisticated signal analyses (Moerel et al. 2012; Giordano et al. 2013; Leaver & Rauschecker 2010), with results sometimes challenging the notion of brain regions dedicated to the abstract encoding of voices (Ogg et al., 2019; Santoro et al., 2017). For instance, regions identified as voice-selective present a response bias to low frequencies typical of voices, even when responding to tones (Moerel et al., 2012), and portions of auditory cortex are sensitive to the degree of harmonic structure present in both artificial sounds and in human vocalizations (Lewis et al., 2009).

The approaches used so far to delineate voice-selectivity in the human brain, mostly relying on functional Magnetic Resonance Imaging (fMRI) or event-related potential recordings with electroencephalography (EEG), present limitations that hinder the characterization of a putative high-level voice-categorization response. For instance, these methods usually imply the subtraction between neural responses elicited by voices and control sounds which occurred at different times, or the regression of parameters linked to low-level properties of sounds, when in fact the subtracted components might be a part of the expression of a response to voices (Frühholz and Belin, 2018).

Here we shed light on the nature of voice-selective responses in the human brain by proposing a new approach to identify these automatic responses objectively and directly (i.e. without subtraction/regression). This approach relies on electroencephalographic recordings and, more specifically, on “EEG frequency tagging” (Regan 1989). Frequency tagging builds on the principles of so-called steady-state evoked responses: under periodic external stimulation, the brain region encoding that input responds at the exact same stimulation frequency (Norcia, Appelbaum, Ales, Cottereau, & Rossion, 2015 for review). We developed a Fast Periodic Auditory Stimulation (FPAS) paradigm adapted from studies in vision, in particular to study face (Retter and Rossion, 2016; Rossion et al., 2015) and letter/word categorization (Lochy et al., 2015). Specifically, participants listened to sequences of heterogeneous sounds presented at a periodic rate of 4 Hz. Critically, each third sound presented in the sequences was a (different) human voice excerpt, so that voices were presented at a periodic rate of 1.333 Hz. In the EEG frequency domain, a response at the sound presentation frequency would reflect shared processes between all sounds, while a putative activity at the voice presentation rate would emerge only if the participant’s brain successfully *discriminates* human voices from other sounds and *generalizes* across all the diverse vocal samples presented (to maintain periodicity). To further assess whether low-level properties alone could elicit voice-selective responses, we included a second stimulation sequence with identical periodicity constraints using the same sounds but frequency-bins scrambled (Dormal et al., 2018) to preserve the overall frequency content of the original sounds while disrupting their harmonicity and intelligibility. In a second experiment, we implemented a sequence presenting voices among musical instrument notes that were matched for pitch, harmonicity-to-noise ratio (HNR) and spectral center of gravity to control for frequency content of the sounds, harmonicity and within category homogeneity (Belin et al., 2011, 2002).

In summary, the goal of the present study was both conceptual and methodological. We aimed to develop a FPAS paradigm combined with EEG to test whether and to which extent voice-selective responses are partially independent from low-level acoustic features. Since FPAS provides a marker for categorization that is objective, direct and does not require overt responses to voices from the participants, we expect our observations to hold significant value for further characterization of the nature of voice-selectivity in the human brain.

## 2. Materials and methods

### 2.1 Experiment 1 – Voice versus object sounds

#### 2.1.1 Participants

EEG was recorded in twenty participants (age range 19-26 years, 10 female) in experiment 1. Data from four participants were excluded due to the presence of EEG artefacts. All participants were right-handed and reported normal or corrected to normal vision, normal hearing and no history of psychiatric or neurological disorders. The experiment was approved by the local ethical committee of the University of Louvain (Project 2016-25); all participants provided written informed consent and received financial compensation for their participation.

#### 2.1.2 Stimuli

Individual sounds used to create the *standard sequences* were 250 ms long, leading to a base stimulation frequency of 4 Hz (1/250ms) and a target frequency of 1.333 Hz (4Hz/3: vocal stimuli presented each third sound) and were selected in an effort to be as heterogeneous and variable as possible. We selected 137 non-vocal stimuli including environmental sounds (e.g. water pouring, rain), musical instruments, sounds produced by manipulable and non-manipulable objects (e.g. telephone ringing, ambulance siren). We selected 55 vocal stimuli including speech and non-speech vocalizations pronounced by speakers of different sex, age and emotional states. Stimuli were extracted from various sources including online databases, extracts from audiobooks and the Montreal Affective Voices dataset (Belin et al., 2008). These stimuli were then frequency scrambled using the method described in Dormal et al. 2018 to create the *scrambled sequences*. Specifically, we applied a fast Fourier transformation to vocal and non-vocal sounds and created frequency bins of 200 Hz. Within each of these frequency windows, we shuffled the magnitude and phase of each Fourier component. We then performed an inverse Fourier transform to the signal and then applied the original sound envelope to the scrambled sound. As a result, the scrambled sounds have frequency content and spectral-temporal structure (change of energy of some frequencies over time) that are almost identical to that of the original stimuli (Figure 1C). However, the harmonicity is altered and the intelligibility of the stimuli is disrupted as confirmed by the behavioral experiment (experiment 3, Figure 5). All sounds were equalized in overall energy (RMS) and faded-in and –out with 10 ms ramps in order to facilitate individual sounds segregation and avoid clicking.

**Figure 1.**
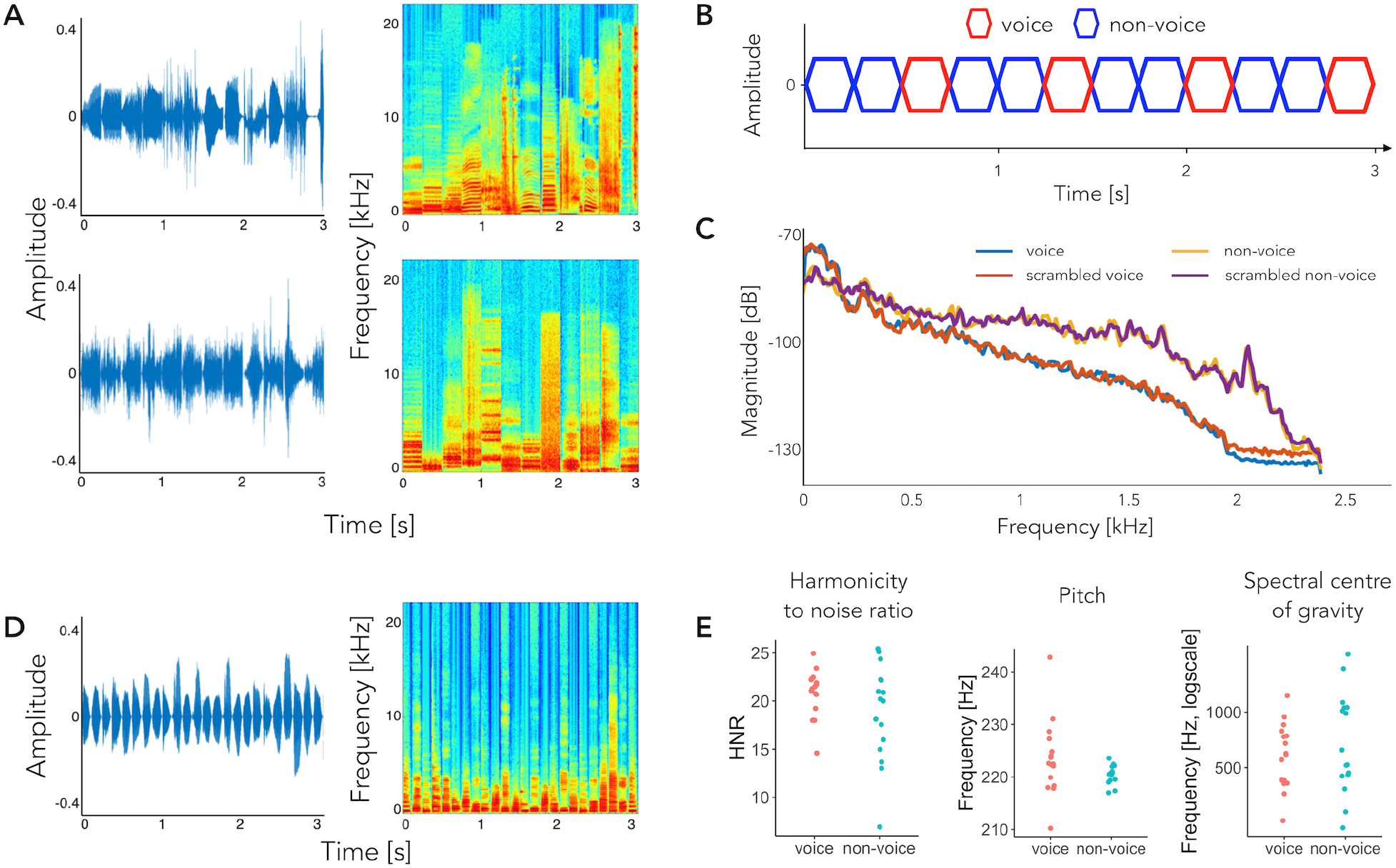
Experimental design for experiments 1 (A, B, C) and 2 (D, E). A) 3 s excerpts of the sequences for the standard (top) and scrambled (bottom) sequences (left) with their relative spectrograms (right). B) Schematic representation of the paradigm. C) Standard and scrambled sequences: Bode magnitude plot expressing the magnitude in decibels as a function of frequency for the averaged sounds of vocal and non-vocal stimuli separately, for standard and scrambled sequences. D) 3 s excerpts of the harmonic sequence (left) with its spectrogram (right). E) Acoustic features as a function of sound category: harmonicity-to-noise ratio, pitch and spectral center of gravity.

#### 2.1.3 Procedure

A schematic representation of the experimental design is shown in Figure 1B. Sounds of the same duration were presented one after another to create periodic auditory sequences. In particular, sounds were presented such that each third sound was a human voice. Vocal and non-vocal samples were selected from various sources and were as heterogeneous as possible to represent the variability characteristic of a sound category (and thus also naturally increasing the variability of low-level acoustic features). No auditory category other than voice was presented periodically. Participants listened to two different sequence types that were created to measure voice selectivity using naturalistic stimuli (*standard sequence*) and to control for low-level acoustic confounds (i.e. frequency content, *scrambled sequences*). If, at the target (voice) rate, the standard and the scrambled sequences evoke responses that are quantitatively (amplitude of the response) and qualitatively (topographical map, pattern of harmonics) similar, it would indicate that we are interpreting as voice-selective responses that are elicited by frequency content alone. Each stimulation sequence was 64 s long, including 2 s of fade-in and -out during which the presentation volume raised gradually from 0 to the maximum at the start of the sequence, and vice versa at the end of the sequence. Fading-in and –out were introduced to avoid abrupt movements that the sudden onset of the sequences could have provoked and that would have introduced artefacts in the data. Sequences for both conditions were presented four times in a pseudorandomized order. Individual sequences with a new pseudo-randomized order of stimulus presentation were generated before testing for each repetition and for each individual participant in an effort to increase generalization; all sequences played during the testing therefore constitute unique exemplars that share presentation parameters (base and target frequencies) and the sounds which were used to build them, but systematically presenting them in different orders in each sequence and each participant. During testing, participants were asked to perform an orthogonal non-periodic task: they had to press a button whenever they heard a sound that was presented at a lower volume as compared to the volume of the other sounds in the sequence. Volume reduction was obtained by decreasing the sounds’ root mean square values of a factor of 12.5. Each sequence contained six attentional targets (including both vocal and non-vocal sounds) that were introduced in a pseudorandomized order (excluding fade-in and -out period at the start of the sequence). Participants were required to listen to the sequences and perform the task blindfolded sitting at 90 cm distance from the speakers.

#### 2.1.4 EEG acquisition

The EEG was recorded with a Biosemi Active Two system (https://www.biosemi.com/products.htm) with 128 Ag-AgCl active electrodes at a sampling rate of 512 Hz. Recording sites included standard 10-20 system locations as well as intermediate positions (position coordinates can be found at: https://www.biosemi.com/headcap.htm). The magnitude of electrode offset, referenced to the common mode sense (CMS), was held below ±50 mV.

#### 2.1.5 Analysis

Data analysis was performed using the Letswave5 toolbox (https://github.com/NOCIONS/Letswave5) and the FieldTrip toolbox (Oostenveld et al., 2011) running on Matlab_R2016b (MathWorks, USA), custom-build scripts in Matlab_R2016b and RStudio.

#### 2.1.6 Pre-processing

A fourth order Butterworth band-pass filter with cut-off values of 0.1-100 Hz was applied to the raw continuous EEG data of each participant. Electrical noise at 50 Hz, 100 Hz and 150 Hz was attenuated with a FFT multi-notch filter with a width of 0.5 Hz. Data were then downsampled to 256 Hz to facilitate data handling and storage. Subsequently, data were segmented into 69 s long epochs (we will use the term epoch in the analysis sections to refer to the EEG data relative to one sequence of stimulation) to include 2 s before the onset of the fade-in and 3 s after the offset of the fade-out of the stimulation sequences. At this stage, after visual inspection of the data, four subjects were excluded from further analysis as their EEG trace was highly contaminated by artefacts. In the remaining sixteen participants, noisy channels were linearly interpolated with the closest neighboring channels (three/four, considering electrodes representing the full area around the interpolated one). This procedure was carried on six participants with no more than 5% of the electrodes (Bottari et al., 2020; Retter et al., 2020) being interpolated in each participant (mean number of channels interpolated considering all sixteen participants is 1, range: 0-6; of the channels interpolated across the six participants, 6 were posterior-occipital electrodes, 2 central parietal, 1 central, 1 temporal, 1 fronto-temporal, 4 fronto-central and 1 anterior-frontal). The same analyses performed on the ten participants on which no channel interpolation was performed and on the entire group (with interpolation) led to very similar results. Epochs that presented multiple artefacts after channel interpolation were removed, with no more than one epoch per condition per participant being excluded. All one hundred twenty-eight EEG channels were then re-referenced to the common average of all electrodes.

#### 2.1.7 Frequency domain analysis

Considering the frequency resolution (1 / duration of the sequence) and the frequency of stimulation (target frequency), pre-processed data were re-segmented so as to contain an integer of voice presentation cycles to avoid overspill of the target-rate response in the frequency domain. Therefore, epochs were re-segmented excluding fade-in and –out and had a final length of 60 s. For each condition and participant separately, epochs were averaged in the time domain to attenuate EEG activity not in phase with the auditory stimulation. A fast Fourier transformation was applied, resulting in amplitude spectra for each channel, condition and subject. Amplitude spectra ranged from 0 to 128 Hz and had a very high resolution of 0.0167 Hz (i.e., 1/60s), thus allowing to isolate responses at the frequencies of interest and their harmonics. Then, we determined the number of significant harmonics at the group level for target and base frequencies: for each condition separately, we computed the grand average across subjects, pooled all channels together and calculated z-scores on these averaged spectra including as baseline 20 surrounding frequency bins (10 bins on each side excluding the immediately adjacent bins, the local minimum and the local maximum; e.g. Retter & Rossion, 2016). Harmonics of the target and base frequencies were considered as significant if their relative z-scores were higher than 2.32 (i.e. p<0.01, 1-tailed, signal > noise). Consecutive significant harmonics were considered, excluding frequencies corresponding to the base-rate responses for the count of target-rate significant harmonics. Responses are represented as topographical head maps summing baseline subtracted amplitude spectra at significant harmonics for target and base frequencies separately, where baseline was calculated as for the calculation of z-scores. Baseline subtraction prior to quantification of the response enables to take into account the fact that different frequency bands are differentially affected by noise in EEG recordings (Luck, 2014), with typically higher noise at low frequencies (below 1 Hz) and in the alpha frequency band (8-13 Hz). We also compared the sum of the baseline subtracted amplitude of the significant harmonics as elicited by the standard and the scrambled sequences (standard > scrambled, Bonferroni correction for the 128 electrodes). To compute this comparison, we considered the highest number of significant harmonics in any of the two conditions (here, *standard*) knowing that including baseline-subtracted activity at non-significant harmonics to compute the overall response is not detrimental (i.e., adding zeroes). The electrodes that were significant for the standard > scrambled comparison were considered to define the voice-selective region-of-interest (ROIvoice).

To assess the robustness of the method to identify voice-selective responses with an even shorter acquisition time, we conducted the same analysis to individuate significant harmonics considering only the responses elicited by the first stimulation sequence for each condition (i.e., one minute of recording).

Lastly, we performed source localization to identify the generators of the voice-selective response. Source localization was implemented here with Dynamic Imaging of Coherent Sources (DICS, Gross et al., 2001) following the method as described in Popov et al., 2018. Using the cross-spectral density (CSD) calculated at the sensor level, DICS estimates the interaction between sources at a particular frequency: in this case, at the voice presentation frequency (1.333 Hz). This beamformer was chosen since it yields lower localization error despite low SNR when compared to other current density measures (Halder et al., 2019). To attenuate brain activity not in phase with the auditory stimulation, all epochs were averaged in the time domain, then across all subjects for standard and scrambled sequences separately; consequently, a Fourier transform was applied to compute the CSD. The forward model was computed with the segmentation of the MRI152 template, based on which a headmodel was generated using the boundary element model (BEM), characterizing the current conduction and propagation properties in surfaces of scalp, skull and brain. Sources were placed in the brain part of the volume conduction headmodel with a resolution of 5 mm. Further, we performed a manual co-registration of the headmodel and Biosemi electrode coordinates using rotation, translation and scaling of the electrodes on the scalp to match our best visual estimate. Due to the imprecise co-registration of the forward model, as we did not have individual participants’ MRI and precise electrode location on the scalp, we did not focus on the exact anatomical areas of the brain generators, but limited our attention to compare the voxel-by-voxel activity (i.e. coherence) between the two conditions. A common spatial filter was computed by appending the data of the two conditions to localize each condition. A regularization parameter of 5% was used. Then, we calculated the difference of coherence values between the standard and scrambled conditions taking the whole brain into consideration to estimate an overall activity at the target frequency. We hypothesized to obtain higher coherence values for the standard condition.

Although source analysis is introduced to suggest a link with previous neuroimaging studies, results should be interpreted cautiously, not only because of the indeterminate nature of source localization from scalp voltage potentials but also because we did not collect MRI scans of individual participants, nor the electrode positions on their scalp during the EEG sessions, important elements that would enhance the precision and accuracy of source localization (Akalin Acar and Makeig, 2013). We therefore consider the results of the source localization for visualization purpose only and the main statistical inferences were done on the scalp data.

### 2.2 Experiment 2 – Voice versus musical instruments controlled for low level acoustic properties

Materials and methods were the same as for experiment 1 unless specified below.

#### 2.2.1 Stimuli

Sounds were extracted from a database provided by Agus and collaborators (Agus et al., 2017; Goto et al., 2003). Stimuli were 128 ms, therefore the base and target frequencies of what we will refer to as *harmonic sequences* were approximately 7.813 Hz and 2.604 Hz, respectively. Vocal stimuli (16 exemplars) consisted in vowels /a/, /e/, /i/ and /o/ sung in the note A3 by two male and two female singers. Non-vocal sounds consisted in sixteen different musical instruments playing the note A3 (oboe, clarinet, bassoon, saxophone, trumpet, trombone, horn, guitar, mandolin, ukulele, harpsichord, piano, marimba, violin, viola, and cello, for a total of 16 stimuli). Importantly, vocal and non-vocal sounds implemented for this sequence type were matched for pitch (M = 223.5 Hz, SD = 7.1Hz for voices, M = 220.6 Hz, SD = 1.85 Hz for instruments, Welch Two Sample t-test t(17.03) = 1.56, p = 0.14), spectral center of gravity (Mann-Whitney Test, W = 107, p = 0.45) and harmonicity to noise ratio (M = 20.9 dB, SD = 2.5 dB for voices, M = 18.9 dB, SD = 5.0 dB for instruments, Welch Two Sample t-test t(21.8) = 1.45, p = 0.16, Figure 1E). Stimuli were equalized in overall energy (RMS) and their RMS value was set as to match the energy of the sounds of experiment 1. Sounds were then faded-in and –out with 10 ms ramps in order to facilitate individual sounds segregation and avoid clicking.

#### 2.2.2 Procedure

*Harmonic sequences* were generated following the same procedure as reported for experiment 1, presenting sounds one after another with each third sound being a voice. Sequences were 65,5 s long including 2 s of fade-in and -out. As in experiment 1, individual sequences were created before testing for each repetition and for each individual participant and included six attentional targets consisting in sounds played at a lower volume. Participants were required to listen to four different harmonic sequences and press a button whenever they perceived a sound played in a lower volume sitting blindfolded at 90 cm from the speakers.

#### 2.2.3 Analysis

Data were analyzed as in experiment 1 with the epochs being re-segmented from 2.048 s after the onset of the sequences (to exclude fade-in and to start segmenting from the onset of the first sound at 100% volume) and had a final length of 59.9 s to contain an integer of voice presentation cycles and avoid overspill of the target-rate response in the frequency domain. The number of significant harmonics for the responses at the target and base frequencies were calculated as in the previous experiment considering all four stimulation sequences or the first sequence alone. We then performed a region-of-interest analysis by averaging the sum of the baseline subtracted amplitude of the significant harmonics of the target of the electrodes of the ROIvoice as defined in experiment 1 and contrasted the resulting activity against 0.

#### 2.3 Experiment 3 - Behavioral detection of voices

To validate that the sounds presented in the two EEG experiments could effectively be categorized as vocal/non-vocal sounds and to assess the effectiveness of the sound scrambling procedure in altering intelligibility, we conducted a behavioral experiment in which participants had to classify all sounds, presented either in short sequences or in isolation, as vocal or non-vocal sounds.

#### 2.3.1 Participants

Sixteen participants (age range 18-26 years, 9 female), four of which had previously participated in the EEG experiments, took part in the behavioral experiment. All participants reported normal or corrected to normal vision, normal hearing and no history of psychiatric or neurological disorders. The experiment was approved by the local ethical committee of the University of Louvain (Project 2016-25); all participants provided written informed consent and received financial compensation for their participation.

#### 2.3.2 Stimuli

Auditory stimuli were the same as for experiments 1 and 2, all equalized so to have the same overall energy (RMS) and faded-in and -out with 10 ms ramps.

#### 2.3.3 Procedure

Participants listened to sounds corresponding to the three sequence types of experiments 1 and 2 (*standard, scrambled* and *harmonic*) that were presented either embedded in short sequences of five sounds (task *sequence*) or in isolation (task *isolation*). For the *sequence* task, short sequences were created so that they could contain either one vocal sound or none (50% / 50% of occurrences). In particular, when a vocal sound was presented in a short sequence, it was inserted as the third sound to mimic the structure of sequences implemented for the electroencephalographic experiments. For each condition, 80 sequences were created (40 with one voice, 40 without voices), for a total of 240 trials (1 short sequence of the behavioral experiment = 1 trial). In the *isolation* task, for the sequence types *standard* and *scrambled* we presented 33 vocal and 67 non-vocal sounds extracted randomly for each participant to reproduce the 1 to 2 ratio of vocal and non-vocal stimuli presented in sequences in the EEG experiment. For the *harmonic* sequence, we presented all 16 vocal and 16 non-vocal stimuli. Sounds from each of the three conditions were presented once in a randomized order, for a total of 132 trials (1 sound = 1 trial). The *sequence* and *isolation* tasks were presented one after another. Each trial consisted in the presentation of one short sequence (*sequence*)/one sound (*isolation*) after which participants had to indicate whether they heard a voice or not with a button press and a response was required in order to initiate the following trial. Participants performed the task blindfolded. The experiment was implemented on Matlab_R2016b (MathWorks, USA) using the Psychophysics Toolbox extensions (Brainard, 1997; Kleiner et al., 2007; Pelli, 1997).

## 3. Results

### 3.1 Experiment 1 – Voice versus object sounds

We expected clear responses at the base frequency (BF, 4 Hz) and harmonics (multiples of BF: 2BF, 3BF, etc.) for both the standard and the scrambled sequences, a response at the base frequency reflecting shared processes between all sounds. Then, we predicted to observe – or not, depending on the sequence type – responses at the target frequency (TF, 1.333Hz) and the harmonics. Critically, a response at the target frequency would arise only if the participant’s brain successfully discriminates human voices from other sounds and generalizes across voices. We predicted that the standard sequences would elicit a voice-selective response reflecting selective responses to vocal versus non-vocal sounds. We also hypothesized that if voice-selective responses evoked by the standard condition were the mere by-product of frequency content, the scrambled condition would have elicited responses quantitatively (in terms of magnitude of the response) and qualitatively (in terms of topographical distribution of the response) similar to the standard condition.

#### 3.1.1 Base frequency

We observed significant responses for the first two harmonics of the base stimulation frequency, i.e. at 4 and 8 Hz, for both the standard and scrambled sequences. The scalp topographies of the two conditions obtained summing baseline corrected amplitudes at the two significant harmonics did not differ, suggesting that individual sounds were similarly processed between the standard and scrambled conditions, with responses peaking over central and occipital electrodes (Figure 2).

**Figure 2.**
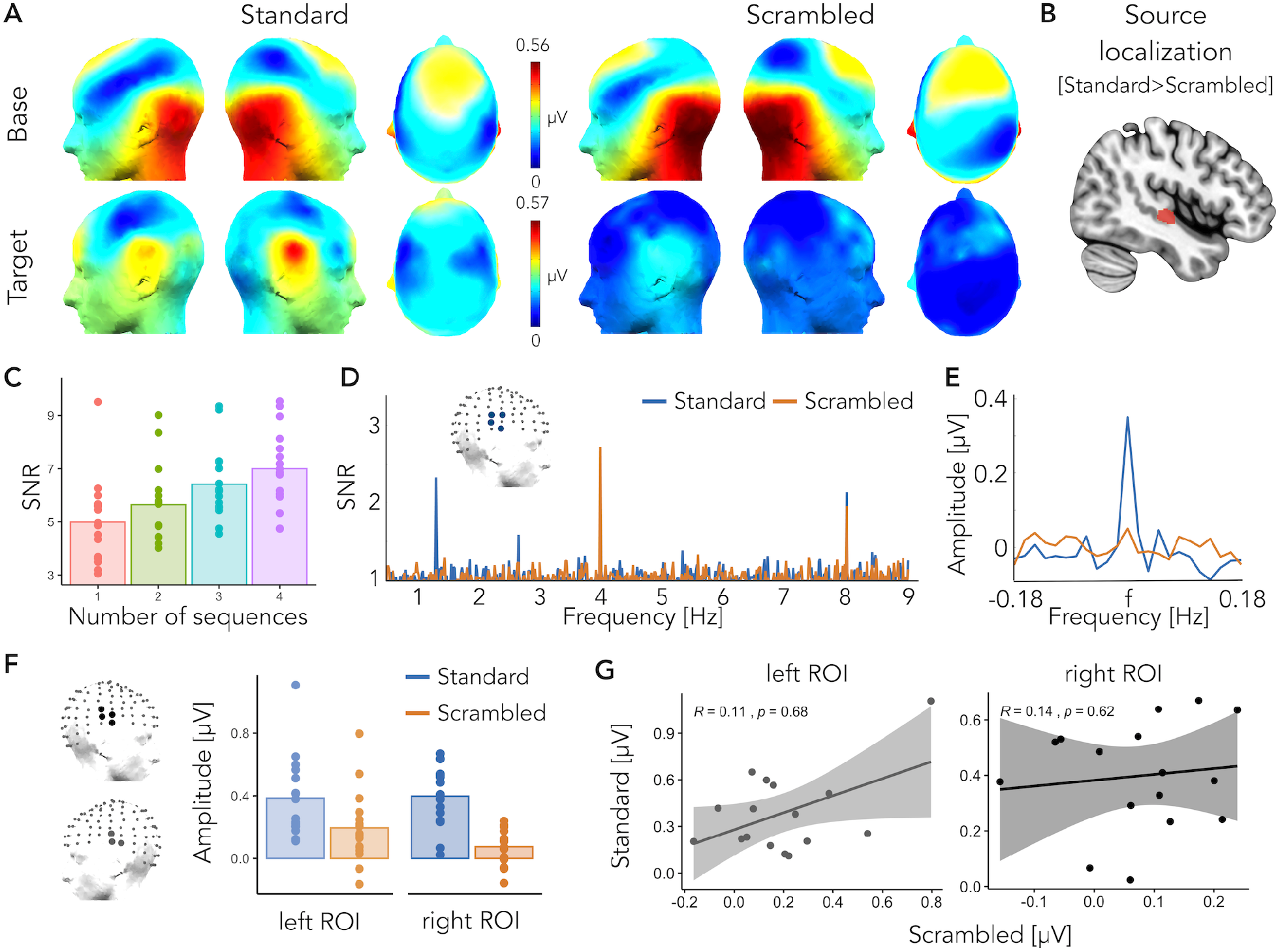
Experiment 1 – Voice versus object sounds. A) Responses at the base (top) and at the target frequency (bottom) for the standard (left) and scrambled (right) conditions as topographic head plots. Topographic head plots were obtained by summing the baseline subtracted amplitude at the significant harmonics. B) Source localization analysis revealed stronger coherence values at the target frequency for the standard condition (standard > scrambled) over right superior and middle temporal areas (voxels with intensity above the 95^th^ percentile are highlighted in red). C) Signal-to-noise ratio (SNR) of voice-selective responses obtained by averaging SNR values of ROIvoice electrodes as a function of number of stimulation sequences/minutes of recording for the standard condition. D) SNR averaged spectra of ROIvoice electrodes. E) Sum of the baseline corrected amplitude (considering 10 bins on each side, excluding the immediately adjacent ones) of voice selective responses at the four significant harmonics for the standard (blue) and scrambled condition (orange). F) Responses at the target frequency for the standard (blue) and scrambled (orange) conditions as the sum of baseline subtracted amplitude at four harmonics in the left and right ROIs. G) There is no correlation between responses at the left and right ROIs across the standard and scrambled conditions.

#### 3.1.2 Target frequency

The standard sequence elicited robust voice-selective responses that were significant at the group level for the first four harmonics of the target frequency (at 1.333 Hz, 2.666 Hz, 5.333 Hz and 6.666 Hz). For the scrambled sequence, we observed a weak but significant response only at the first harmonic of the target frequency (at 1.333 Hz). We quantified voice-selective responses as the sum of the baseline subtracted amplitudes of the highest number of significant harmonics in any of the two conditions (here: standard, four harmonics) knowing that including baseline-subtracted activity at non-significant harmonics to compute the overall response is not detrimental (i.e., adding zeroes; Retter et al., 2018). For the standard condition, voice-selective responses peaked bilaterally at superior temporal electrodes (Figure 2): TP8h, CP6, T8h, T8 on the right hemisphere and T7h, T7, TP7 on the left hemisphere (contrasting the response against zero, one-sided t-test, reported p-values are Bonferroni corrected, Cohen’s d values are reported referred to as “d”, TP8h: t(15) = 6.68, p = 0.0005, d = 1.67; CP6: t(15) = 5.51, p = 0.0038, d = 1.38; T8h: t(15) = 6.89, p = 0.0003, d = 1.72; T8: t(15) = 7.16, p = 0.0002, d = 1.79; T7h: t(15) = 5.62, p = 0.0031, d = 1.40; T7: t(15) = 5.44, p = 0.0043, d = 1.36; TP7: t(15) = 5.56, p = 0.0035, d = 1.39), the topography of the response being consistent across participants (Figure 3). No electrode reached significance when we computed the same contrast for the scrambled condition. Overall, in regions where the response was peaking for the standard condition, the amplitude of the response to voice-scrambled stimuli was of 50.6% of the response to voices for the left ROI (T7h, T7, TP7) and only 18.8% for the right ROI (TP8h, CP6, T8h, T8; Figure 2F). Moreover, the magnitude of the responses to the standard and scrambled conditions over these two region-of-interests did not correlate across participants (Spearman’s rho correlation, left ROI: r_s_ = 0.11, p = 0.68, right ROI: r_s_ = 0.14, p = 0.62; Figure 2G). This suggests that, although scrambled voices could elicit a response, frequency content alone was not enough to elicit a voice-selective response as recorded with the standard sequence. We further investigated that by comparing the responses elicited at the target frequency by the standard and the scrambled sequences; although one of the advantages of the FPAS paradigm is that it does not require a direct comparison between conditions, we decided to include this extra step as a proof of principle since we introduce oddball fast periodic stimulation with high-level, naturalistic sounds for the first time here. First, for each condition, participant and electrode separately we summed the responses of the highest number (i.e., four) of significant harmonics in any of the two conditions. Then, comparing the responses for each participant and electrode (standard > scrambled, paired-sample t-test, Bonferroni corrected), we isolated four significant superior temporal electrodes over the right hemisphere: TP8h, CP6, C6 and T8 (TP8h: t(15) = 6.21, p = 0.0011, d = 1.55; CP6: t(15) = 5.02, p = 0.0097, d = 1.26; C6: t(15) = 4.71, p = 0.0179, d = 1.18; T8: t(15) = 4.60, p = 0.0223, d = 1.15, Bonferroni corrected). Non-parametric testing (Winkler et al., 2014) of the standard > scrambled comparison led to very similar results and the choice of a one-sided comparison was made with the a priori assumption that the scrambled condition would have elicited a target response as high - if voice responses could have been explained by frequency content alone - or smaller than the one elicited by the standard sequence. Finally, a region-of-interest was defined considering the four electrodes identified as above (ROIvoice; Figure 2).

**Figure 3.**
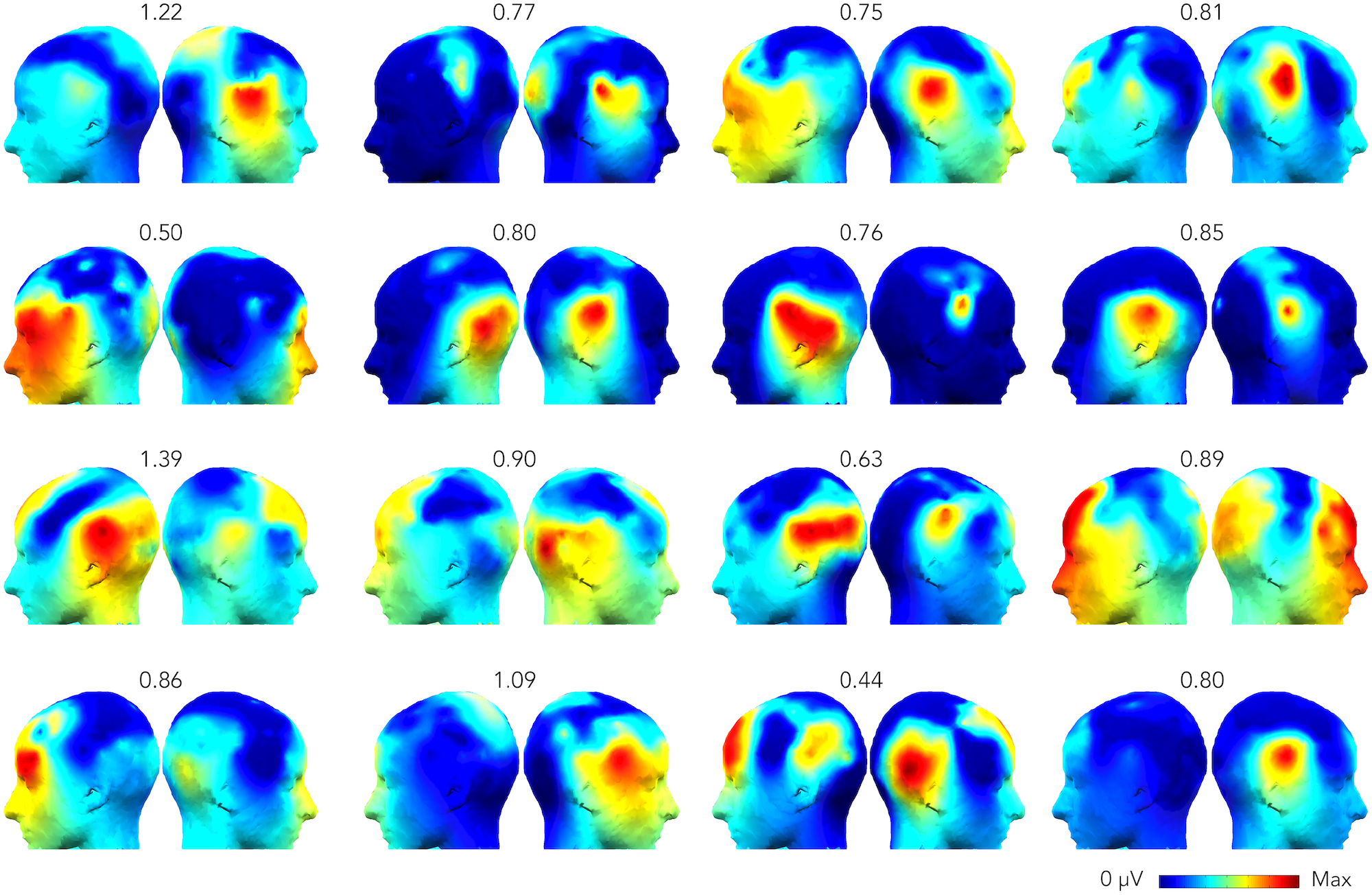
Experiment 1 - individual voice-selective responses. Responses at the voice presentation frequency for each individual. Topographic head plots represent the sum of the baseline subtracted amplitude of significant harmonics as identified at the group level. The scale of each head plot ranges from 0 μV to the maximum (reported on top) for each subject.

To assess whether FPAS was a suitable tool to investigate voice-selectivity at the individual level, we calculated whether these responses were significant in every subject (Liu-Shuang et al., 2016). For each participant, we considered the amplitude spectrum and we pooled together all channels. We chunked epochs that were centered around the harmonics of the target frequency that were significant at the group level (1.333 Hz, 2.666 Hz, 5.333 Hz and 6.666 Hz) and that contained 11 frequency bins on each side. We then summed these epochs together and computed the z-scores at the target frequency considering as baseline the 20 surrounding frequency bins (10 on each side, excluding the immediately adjacent bin). Considering as significant responses whose relative z-scores was higher than 1.64 (i.e. p<0.05, 1-tailed, signal > noise), we were able to find significant voice-selective responses in 13 out of 16 participants (with only 4 minutes of recordings).

To assess the robustness of the method with an even shorter acquisition time, we then performed the same analytical steps done to calculate the number of significant harmonics at the group level considering the first sequence of recording only. For the standard sequence, we were able to identify voice-selective responses that were significant at the group level even with one minute of recording only (1 significant harmonic: 1.333 Hz). The signal-to-noise ratio (SNR) of the response at right superior temporal electrodes (ROIvoice) as a function of number of stimulation sequences (i.e., minutes of recordings, as each sequence is one minute long) is presented for visualization purposes in Figure 2C.

A stronger voice-selective response in the standard condition was observed when compared to the scrambled condition at the source level as well. After implementing source localization, the final voxel coherence values were compared for the two conditions (standard > scrambled) and interpolated to the MRI152 template. Then, the voxels with intensity above the 95^th^ percentile were selected, as illustrated in figure 2B. The maximum value obtained of 0.32 (min-max range: [0 1]) indicates voxel preference for voice presentation frequency (i.e. 1.333 Hz) for the standard over the scrambled sequence. Despite the imprecise source localization due to the limitations outlined in the method section (lack of individual coregistration between electrodes position and brain anatomy), the position of the source reconstructed effect of interest (Figure 2B) lies in the vicinity of the known location of TVAs (Belin et al., 2002, 2000) and are in line with results showing that right anterior STS regions respond more strongly to non-speech vocal sounds than their scrambled versions (Belin et al., 2002).

### 3.2 Experiment 2 – Voice versus musical instruments controlled for low level acoustic properties

We expected to observe a response at the base presentation frequency (7.813 Hz) and its harmonics, this response reflecting processes that are shared among vocal and non-vocal sounds. More crucially, we predicted that, if voice-selectivity is not solely driven by harmonicity and/or pitch of the sounds, we would observe a target-rate response to the harmonic sequences as in these sequences vocal and non-vocal sounds did not differ for these low-level acoustic features.

#### 3.2.1 Base frequency

There was a significant base-rate response for the first two harmonics: at 7.813 Hz and 15.625 Hz, with responses peaking over central and occipital electrodes (Figure 4). The topography of the response was qualitatively similar to the base-rate responses obtained in experiment 1.

**Figure 4.**
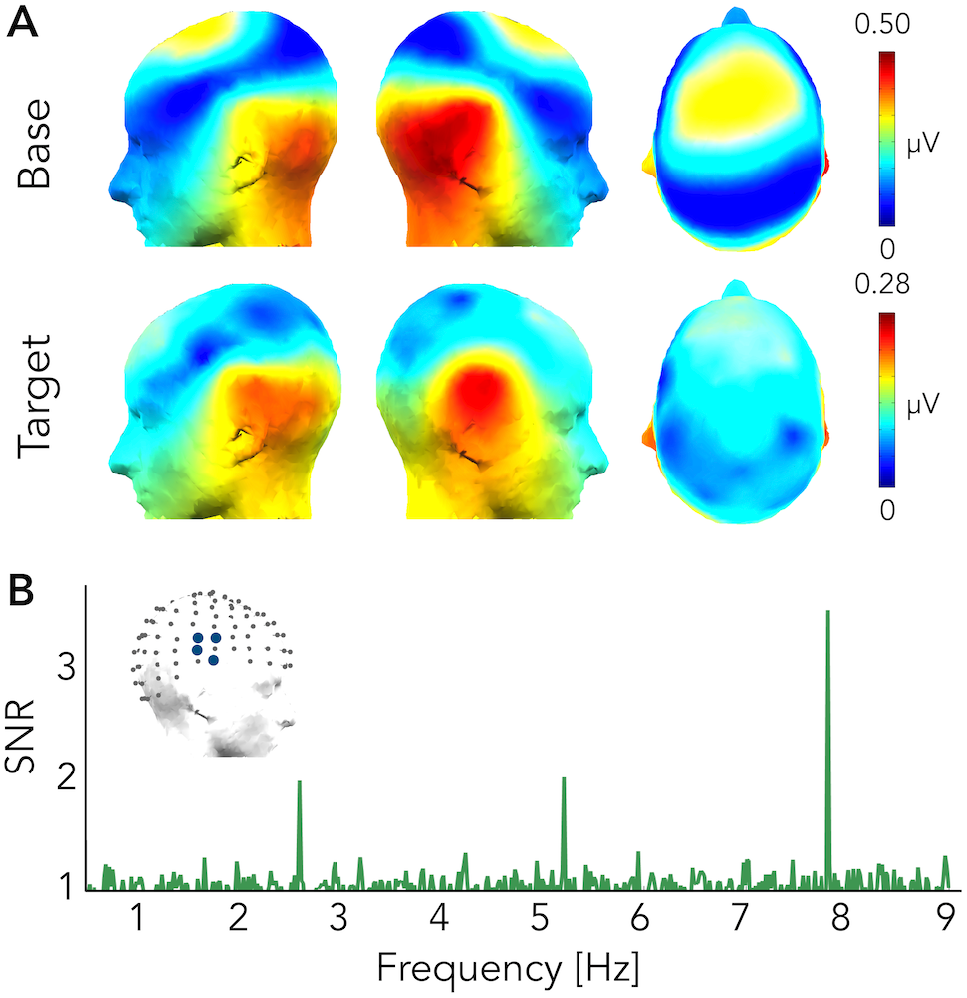
Experiment 2 – Voice versus musical instruments. A) Responses at the base and at the target frequency as topographic head plots. Topographic head plots were obtained by summing the baseline subtracted amplitude at the significant harmonics. B) SNR averaged spectra of ROIvoice electrodes.

**Figure 5.**
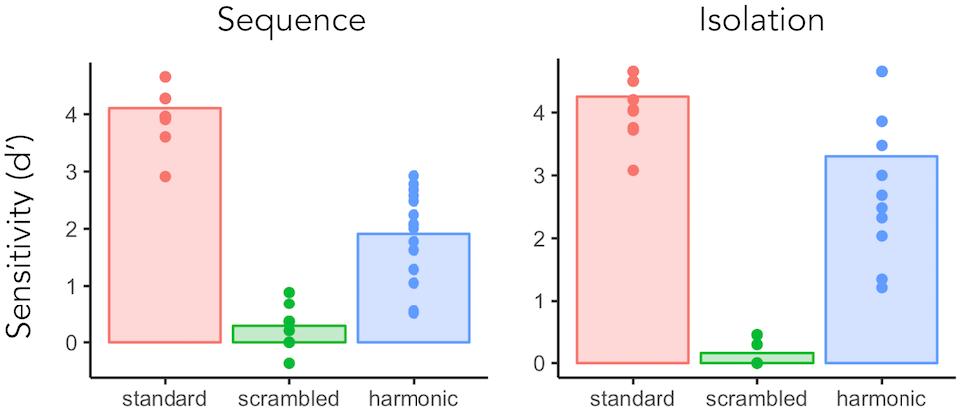
Experiment 3 - Behavioral detection of voices. Sensitivity indices for the sequence (left) and isolation (right) tasks, for the three conditions. The bar plots represent the group average of d’, each dot represents an individual participant’s d’ score.

#### 3.2.2 Target frequency

We observed a significant voice-selective response for the first two harmonics (2.604 Hz and 5.208 Hz). Voice-selective responses peaked over central and superior temporal electrodes bilaterally, the scalp distribution of the response being highly similar to the voice-selective response found in the standard sequence (Figure 4) and being reliable across participants. We then performed a region-of-interest analysis considering the right superior temporal electrodes (ROIvoice) as defined in experiment 1. Baseline subtracted amplitudes at the two significant harmonics were summed for each electrode individually and the resulting responses at the electrodes of the ROIvoice were then averaged together for every participant. One data point was excluded as it deviated more than three standard deviations from the mean of the group and responses were significant against zero (M = 0.152 μV, SD = 0.087 μV, t(14) = 6.78, p = 4.436 × 10^−6^, d = 1.75, 1-tailed t-test).

Voice-selective responses were already present after one minute of recording, showing the resilience of FPAS even with very short acquisition time (one harmonic significant, the second: 5.208 Hz, with two minutes of recording the first two harmonics reached significance at the group level; Figure 4).

As for the standard condition, we then assessed significance of voice-selective responses at the individual level with the same procedure. 15 out of 16 participants showed a significant response (z-score higher than 1.64, i.e. p<0.05, 1-tailed, signal > noise).

### 3.3 Experiment 3 - Behavioral detection of voices

Responses to the *sequence* and *isolation* tasks were analyzed according to signal detection theory. Specifically, sensitivity indices related to the participants’ ability to detect a voice when present was assessed using d-prime (Figure 5). D-prime (d’) constitutes an unbiased quantification of performance in detection tasks as it takes into account both hits and false alarms (Macmillan and Creelman, 2004; Tanner and Swets, 1954). Data points were considered as outliers when deviating from the mean of the group of more (or less) than three times the standard deviation of the group in at least one condition/task, leading to the exclusion of one participant. We first checked whether performance was above chance (d’ = 0) for each condition and for the *sequence* and the *isolation* task separately. Participants performed above chance for all conditions in the sequence (t-test, 1-tail, standard: d’ values mean M = 4.11, SD = 0.45, t(14) = 35.30, p = 2.2 × 10^−15^, Cohen’s d = 9.11; scrambled: M = 0.29, SD = 0.30, t(14) = 3.75, p = 0.001, d = 0.97; harmonic: M = 1.91, SD = 0.77, t(14) = 9.58, p = 7.8 × 10^−8^, d = 2.47) and in the isolation task (t-test, 1-tail, standard: M = 4.25, SD = 0.46, t(14) = 35.71, p = 1.8 × 10^−15^, d = 9.21; scrambled: M = 0.16, SD = 0.21, t(14) = 2.97, p = 0.005, d = 0.77; harmonic: M = 3.30, SD = 1.24, t(14) = 10.29, p = 3.3 × 10^−8^, d = 2.66). Statistical comparisons of conditions were then performed separately for the two tasks using repeated measures ANOVAs. Whenever Mauchly’s test indicated that the assumption of sphericity had been violated, we applied a Greenhouse-Geisser correction to the degrees of freedom. For the *sequence* task, we found a significant effect of condition (F(2,28) = 207.39, p = 1.6 × 10^−17^, η^2^ = 0.898) and post-hoc pairwise comparisons showed that all conditions differed one from another (standard vs. scrambled, p < 2 × 10^−16^; standard vs. harmonic, p = 7.1 × 10^−9^; scrambled vs. harmonic, p = 1.5 × 10^−6^; Bonferroni corrected). The same pattern of performance was found when sounds were presented in isolation (F(1.20, 16.86) = 134.23, p = 6.8 × 10^−10^, η^2^ =0.845; pairwise comparisons: standard vs. scrambled, p < 2 × 10^−16^; standard vs. harmonic, p = 0.038; scrambled vs. harmonic, p = 2.7 × 10^−7^; Bonferroni corrected). Although performance was above chance for the scrambled sounds, d-prime scores were significantly lower than the one achieved in the standard condition (Figure 5), suggesting that our frequency scrambling effectively disrupted the intelligibility of the sounds.

## 4. Discussion

Similarly to the issue of category-selectivity in human visual cortex (Bracci et al., 2017; Peelen and Downing, 2017; Rice et al., 2014), studies have attempted to determine whether category-selectivity in auditory cortices is mostly driven by a biased tuning toward low-level acoustic features or if categorization responses go beyond the acoustic properties of sound and therefore represent a more abstract representation of voices (Giordano et al., 2013; Leaver and Rauschecker, 2010). In fact, as sounds tend to be acoustically similar within an auditory category and dissimilar between categories, differences in cortical responses could be a mere reflection of different acoustic properties across categories of sounds (Ogg et al., 2019; Staeren et al., 2009). One approach to address this question is to test for low-level featural coding. However, although studies using artificial stimuli allow for a careful control of low-level acoustic properties (Lewis et al., 2009; Patterson et al., 2002; Warren et al., 2005), they usually lack ecological validity and underestimate the issue of stimulus-driven response correlation (Norman-Haignere and McDermott, 2018). Moreover, artificial stimuli may fail in eliciting brain activation in a (behaviorally) relevant way, as evidenced by a study revealing different tonotopic maps obtained with natural sounds and pure tones (Moerel et al., 2012).

In this study, we developed a Fast Periodic Auditory Stimulation (FPAS) paradigm as a powerful mean to investigate voice-selectivity in the brain with the aim of disentangling between a categorization response to voices and the contribution of low-level acoustic features that are typical of voices (e.g. high harmonicity, characteristic frequency ranges, specific change of energy over time) with EEG. The frequency constraint of this approach allowed us to individuate robust voice-selective responses objectively – at known stimulation frequencies – and automatically: not only participants did not have to overtly respond to voices, thus avoiding the contamination of the response from attentional and decisional processes (Levy et al., 2003; Von Kriegstein et al., 2003), but also no subtraction between responses elicited by different auditory categories was required. That is, while traditional M/EEG approaches investigate whether there are differences in isolated responses elicited by different classes of stimuli (Charest et al., 2009; De Lucia et al., 2010; Levy et al., 2003; Murray et al., 2006), in our paradigm this differentiation is implicit. Moreover, the relatively low-speed presentation of stimuli required by many traditional experimental paradigms preclude from the possibility of presenting a high number of stimuli that could be representative enough of an auditory category and therefore responses observed to such a subset of voice samples might not reflect a *generalized* response to voices (Giordano et al. 2013). In addition, low-speed presentation of stimuli might fail in fully characterising voice processes as they occur in daily-life, since we are experts in extracting voice features almost effortlessly in highly dynamic acoustically changing environments.

For a voice-specific response to be captured with the FPAS paradigm, two processes need to occur: the brain has to concurrently *discriminate* voices from non-vocal sounds and to *generalize* this selective response across diverse vocal samples for a response at the target (i.e. voice presentation) rate to occur. FPAS therefore not only allows to characterize a general voice-selective response, as it is elicited by heterogeneous vocal samples and not from a specific subset of those, but also not to cancel out processes elicited by voices that are shared by other sound categories. This can be accomplished in a very short acquisition time (i.e. four minutes or even less) and with a high SNR (Figure 2C, e.g., up to 7, or 600% of amplitude increase with 4 stimulation sequences).

To isolate voice-selective responses that could not be merely explained by low-level acoustic features, we implemented two EEG experiments using the FPAS principle. In the first experiment, participants were presented with two different types of FPAS sequences. In the standard sequence, vocal and non-vocal samples were selected to be as heterogeneous as possible to represent the high variability of sounds of a given category (e.g., voices from speakers from different ages, sex, emotional state with speech and non-speech vocalizations) as encountered in natural environment, as well as to minimize the potential contributions of low-level acoustic features in the voice-selective response (controlling by variability). Specifically, low level acoustic properties would have to vary periodically to participate in the target-specific responses, and the high variability of the voice samples adopted should prevent that. These sounds were then scrambled using small frequency bins (after FFT) in order to generate a sequence in which scrambled sounds had similar frequency content as the original stimuli (Figure 1C) but were not recognizable anymore (Figure 5). We measured robust voice-selective responses at superior temporal electrodes that were significant in the vast majority of the participants with only 4 minutes of recording (Figure 2). Moreover, these responses remained significant at the group level when restricting the analysis to the first minute of recording only, highlighting the robustness of the method even with an extremely short acquisition time. Voice-selective responses could not have been explained by frequency content alone: although one of the advantages of FPAS is that it does not require an explicit comparison across conditions, here, as an extra proof of principle, we compared the responses elicited by the standard and scrambled sequences, highlighting a response expressing over right superior temporal electrodes. This preference also emerged at the source space: although source localization as implemented in our study presents limitations that could hinder the accuracy of our results (see methods section), we localized a standard > scrambled preference over the right superior temporal gyrus and sulci. The observed regions lie in the proximity of the TVAs and are in line with results showing that the right anterior STS regions respond more strongly to non-speech vocal sounds than their scrambled versions (Belin et al., 2002).

It has to be noted that the scrambled condition elicited a weak but yet significant response at the (scrambled) voice presentation rates: we think there might be two - not mutually exclusive-explanations for this weak response. First, this response might be due to the low-level acoustic features shared by voices and scrambled voices (i.e. frequency content): preferential response biases towards acoustic features peculiar of voices have been observed in voice-selective regions (Moerel et al., 2012) and, despite the fact that voice-selective responses elicited by the standard condition could not be well explained by low-level acoustic features alone, these features may nevertheless weakly contribute to the response. Second, although the scrambling procedure disrupted the intelligibility of the sounds, with original sounds being more accurately recognized as voices or not than scrambled sounds, participants’ performance in categorizing scrambled sounds was above chance level, suggesting that some residual sound recognizability might have still been present.

To further investigate the nature of voice-selective responses, we designed a second FPAS experiment using sequences built from sung vowels and musical instruments that were matched in terms of pitch, spectral center of gravity and harmonicity to noise ratio. The observation of robust voice-selective responses despite controlling for the above-mentioned acoustic features speaks in favor of a categorization response to voices that is at least partially independent from some of the most intrinsic acoustic features of voices. The similar scalp topographies elicited by the standard and harmonic sequences suggest a similarity between the selective responses to voices elicited in the two conditions. However, the voice-selective response in the standard sequences was larger in amplitude as compared to the response of the harmonic sequence. Different factors might account for this difference. First, the higher amplitude of response observed in the standard sequence could be due to voices being more easily discriminated from non-vocal sounds, as pointed out by the behavioral data. In fact, even if an overt response to voices is not necessary to elicit a target rate specific response, the easiness at which a voice is perceived could have impacted the amplitude of brain responses. Second, while vocal stimuli of the harmonic sequence were accurately chosen to be matched in acoustic properties with the sounds of musical instruments, it could be argued that those vocal stimuli represent a subset of voices, and thus that the strongest response in the standard condition reflects the response to a more heterogeneous and representative sample of voices of that condition. The choice of the frequencies of stimulation could also have impacted on the magnitude of the responses (Ding et al., 2006; Regan, 1966; Retter et al., 2020): while all parameters proved effective in eliciting target-rate responses, further studies would be needed to indicate which parameters are optimal to capture voice-selective responses, as addressed in the case of the implementation of the frequency tagging approach in high-level vision (e.g., Retter et al., 2020; Alonso-Prieto et al., 2013). Finally, systematic differences in low-level acoustic features between vocal and non-vocal sound potentially present in the standard sequence could have boosted the categorization response to voices. In fact, feature dependence is not in conflict with a categorical coding hypothesis (Bracci et al., 2017), and preferential response biases observed in voice regions to specific acoustic features such as low frequencies (Moerel et al., 2012), harmonic structure (Lewis et al., 2009) and spectrotemporal modulations (Santoro et al., 2017), could serve as a scaffold or facilitate a higher level categorical responses to voices.

In summary, we objectively defined human voice-selective responses independent from low-level acoustic cues that are characteristic of voices with high SNR and in a very short acquisition time using an original Fast Periodic Auditory Stimulation (FPAS) approach. Although it is possible that other acoustic features that were not explicitly controlled for might be at the origin of the recorded responses, the nature of the FPAS design makes it unlikely, as said features should be systematically present in all voices and absent in non-vocal sounds. In other words, activity in voice-selective regions could not be only, or even substantially, accounted for by any basic acoustic parameter tested.

While the goal of the current study was to investigate the existence of a voice-selective response partially independent from some acoustic features, the FPAS paradigm we developed could be a valuable tool for the study of auditory perceptual categorization in the human brain, extending to other categories. Moreover, the high SNR of this technique achievable in a very short acquisition time (i.e., significant responses were observed with one minute of recording) and the fact that no overt response to specific stimuli is required, makes this approach highly promising for studying voice perception in children and clinical populations. Our findings therefore advance our understanding of voice-selectivity in the brain and our method provides an essential foundation for understanding its development in typical and atypical populations.

## Acknowledgments

The authors are grateful to Professor Trevor Agus and Professor Daniel Pressnitzer for sharing the stimuli used in Agus et al. 2017. We would also like to extend our gratitude to Mohamed Rezk and Joan Liu-Shuang for their precious help with the data analysis.

